# Defective Homologous Recombination and Genomic Instability Predict Increased Responsiveness to Carbon Ion Radiotherapy in Pancreatic Cancer

**DOI:** 10.1101/2024.04.08.588534

**Authors:** Brock J. Sishc, Janapriya Saha, Elizabeth Polsdofer, Lianghao Ding, Huiming Lu, Shih-Ya Wang, Katy L. Swancutt, James H. Nicholson, Angelica Facoetti, Arnold Pompos, Mario Ciocca, Todd A. Aguilera, Michael D. Story, Anthony J. Davis

**Author notes:** Current address: Mayo Clinic Florida, Jacksonville, FL, USA.

## Abstract

Pancreatic ductal adenocarcinoma (PDAC) is inherently resistant to conventional chemo-and radiation-therapy. However, clinical trials showed that carbon ion radiotherapy (CIRT) with concurrent gemcitabine can be effective for treating unresectable locally advanced PDAC. In this study, we aimed to determine features that could identify patients who would benefit most from CIRT. A panel of human PDAC cell lines with various genetic backgrounds was leveraged to determine whether a subset could be identified that preferentially responds to CIRT. The cell lines displayed a differential response to CIRT as compared to γ-rays as measured by relative biological effectiveness (RBE) calculated at 10% survival, which ranged from 1.96 to 3.04. Increased radiosensitivity correlated with decreased DNA double strand break (DSB) repair as measured by γH2AX foci resolution. We determined that the cell lines most sensitive to CIRT are defective in the homologous recombination (HR) DSB repair pathway and/or have high genomic instability due to elevated replication stress. Next, this knowledge was utilized to assess whether the HR pathway could be targeted to potentiate CIRT *in vitro*. It was determined that pretreating a radioresistant PDAC cell line with the HR inhibitor, B02, resulted in a marked increase in sensitivity to CIRT when treated with high linear energy transfer (LET) radiation in the spread-out Bragg peak (74.1-89.3 keV/μm) but not at the entry LET (13.0-16.4 keV/μm) *in vitro* as opposed to that seen with a NHEJ inhibitor. These data suggest a greater therapeutic index with the combination therapy in the tumor over normal tissues based on LET distribution. These data support the notion that PDAC tumors with defects in HR and/or those with high inherent replication stress respond to CIRT without the concern for excessive normal tissue toxicity. With the advent of agents targeting HR, the difference in tumor cell response in the entry region vs within the SOBP based upon the respective LETs of those regions of the beam profile, will be superior to agents that target NHEJ.

## INTRODUCTION

Pancreatic ductal adenocarcinoma (PDAC) is the fourth leading cause of cancer death in both men and women in the United States (U.S.)^1^. The primary curative treatment is surgery for resectable disease, however only 15-20% of PDAC patients are eligible for this treatment option due to the location of the tumor and disease stage^2^. Gemcitabine (GEM) is the only active single agent chemotherapeutic that improves survival, however, combination chemotherapy with either folinic acid, fluouracil, irinotecan, oxaliplatin (FOLFIRINOX) or albumin-bound paclitaxel (nab-PTX)/GEM resulted in greater survival impact for localized and metastatic disease PDAC^3–5^. Thus, these combination chemotherapies are now considered standard of care. Clinical studies have shown that a combination of chemotherapy and radiation can transition an unresectable PDAC to a resectable state in roughly 20% of cases. However, the role of radiotherapy in improving survival in localized disease remains to be demonstrated, likely due to inherent tumor radioresistance, lack of biological triage, and/or an insufficient total dose of ionizing radiation (IR)^6^. Despite improvements in treatment options, the 5-year overall survival remains low at just over 10%^7^. Thus, there is a great need for the development and validation of additional innovative, potent, and targeted/selective treatment approaches.

Heavy ion therapy, such as carbon ion radiotherapy (CIRT), could be a game changer for PDAC. Particle radiotherapy in general, and CIRT in particular encompass numerous physical and biological therapeutic advantages when compared to conventional radiotherapy and some chemotherapeutics. The physical advantages include the generation of a spread-out Bragg peak (SOBP) to focus the most damaging portion of a particle track inside the tumor, an enhanced dose distribution that more effectively spares nearby, at risk structures, lateral beam focusing to improve field homogeneity, dose verification allowing for real-time treatment modification, an increased linear energy transfer (LET) that causes more damage to the tumor cells, and finally the capability to magnetically steer the ion beam leading to a more precise dose delivery^8^. The biological advantages of CIRT include a higher relative biological effectiveness (RBE) damaging cells more effectively per unit of physical dose as compared to photons or protons, a reduced dependency on molecular oxygen, which results in a lower oxygen enhancement ratio (OER) thus creating more damage in a hypoxic tumor, and the generation of complex DNA damage which leads to more persistent stress and cell death^8,9^. Furthermore, a carbon ion beam offers an ideal energy distribution which induces a maximum ionizing effect at the site of the tumor and less damage to the surrounding normal tissue, leading to the prediction that CIRT will result in better tumor control with fewer side effects than conventional radiotherapy.

Clinical data has been encouraging; as a Phase I dose escalation study in unresectable PDAC by the Japanese Working Group for Pancreatic Cancer showed a 2-year survival rate of 48% for patients treated with 45.6-55.2 GyE and concurrent gemcitabine^10^. A follow up single institution study demonstrated a 53% 2-year survival for patients that received 55.2 GyE^11^. Local control in this latter group was 82%, suggesting that CIRT is improving local control of later stage disease that is not observed with conventional or hypofractionated X-ray radiotherapy^6,11,12^.

Due to the concentrated energy deposition within a confined volume, carbon ions cause DNA damage of greater complexity. A special feature of this densely ionizing radiation is the induction of clustered DNA lesions, which is defined as two or more DNA lesions, such as DNA double strand breaks (DSBs), single strand breaks, or base damage, within one or two helical turns of the DNA^13,14^. Multiple studies have revealed that as LET increases, DNA repair slows as complex DNA lesions are more difficult to repair^15–17^. In response to clustered DNA damage, cells activate multiple DNA damage response (DDR) pathways. The most toxic of the CIRT-induced DNA lesions are DSBs, which if left unrepaired or are misrepaired can result in genomic instability or cell death by a number of mechanisms including apoptosis, mitotic catastrophe, or senescence. DSBs are repaired by three pathways: homologous recombination (HR), non-homologous end joining (NHEJ), and alternative end joining (alt-EJ)^18^. Multiple studies have addressed the different contributions of the NHEJ and HR pathways to DSB repair according to the complexity of the DSBs generated, with HR being more important for the repair of high-LET than low-LET generated DNA lesions^19–21^. However, the contributions of the NHEJ and HR pathways to the repair of clinical carbon ion beam-induced DSBs have not been clarified in human PDAC cell lines.

Sequencing and chromosomal copy number variation analyses of PDAC revealed a complex genomic landscape^22,23^. Activating mutations of *KRAS* are near ubiquitous and inactivation of *TP53*, *SMAD4,* and *CDKN2A* occur at rates of >50%. Genomic classification via patterns of variation in chromosome structure identified four subtypes of PDAC that were termed (i) stable, (ii) locally arranged, (iii) scattered, and (iv) unstable^23^. The unstable subtype accounts for 15-20% of human PDACs and the majority of these tumors harbor a mutation(s) in a gene required for the DDR, including *BRCA1*, *BRCA2, PALB2,* and *ATM*. Similarly, mutations in DDR genes are commonly found in inherited forms of PDAC^24,25^. DDR deficiency renders some tumors preferentially sensitive to DNA-damaging agents such as platinum agents and PARP inhibitors^26–28^. Additionally, DDR deficiency has been observed to be predictive for FOLFIRINOX (platinum-containing regimen) efficacy in PDAC^29^. Unfortunately, FOLFIRINOX is suitable only for patients with good performance status given the toxicity. Moreover, the burden of morbidity, and even mortality, associated with platinum chemotherapy is a major challenge complicating the use of this chemotherapeutic regimen ^29^. Therefore, it is hypothesized that a less toxic, tailored and targeted therapy for PDACs with either somatic or germline mutations in DDR genes could be leveraged to improve treatment outcomes and the overall toxicity profile.

In this study, molecular mechanisms of CIRT were identified that impact the therapeutic response in PDAC cell lines. We investigated the contributions of the NHEJ and HR pathways to the repair of carbon ion-generated DNA lesions with the goal of identifying genomic features that can guide the treatment of pancreatic cancer, and distinguish patients who would benefit the most from CIRT. We demonstrate that PDAC cell lines are more sensitive to CIRT than γ-rays and that this correlates with the burden of unrepaired DSBs. Furthermore, we found that the most sensitive cell lines are those with defects in the DDR. If we target a radioresistant PDAC cell line with the HR inhibitor, B02, there is an increased sensitivity to CIRT at the spread out Bragg peak but not at the entry of the depth-dose distribution *in vitro,* indicating an effect potentially driven by LET. Collectively, our data support the notion that PDAC patients or tumors with defects in DDR can benefit the greatest from CIRT. However, if a DDR defect is not identified then the HR or ATR pathways become critical targets to enhance response without complication of adverse normal tissue events.

## RESULTS

### Differential Response of PDAC cell lines to **γ**-ray and ^12^C ion irradiation

PDAC has been shown to be effectively targeted by CIRT clinically^6,10,12^; however, it is unknown whether genetics affect the response to this treatment modality. To assess the impact of genetic variability, we examined the radiation response of a panel of seven human tumor-derived PDAC cell lines with varied genetic backgrounds to both γ-rays and CIRT. Each cell line contained a *KRAS* G12 mutation and additional mutation or deletion events in at least two of the other three commonly mutated genes (*TP53*, *CDKN2A*, and *SMAD4)* found in PDAC (**Supplementary** Fig. 1A). Furthermore, we noted various mutations of known or unknown significance in DNA damage response (DDR) genes and other putative oncogenic mutations (**Supplementary** Fig. 1B). Clonogenic survival curves were generated using^137^Cs for γ-ray (low LET) irradiation at UT Southwestern Medical Center (UTSW) in Dallas, TX and carbon ions (high LET) irradiation at the National Center for Oncological Radiotherapy (CNAO) in Pavia, Italy. As demonstrated in **Fig. 1A and B**, all cell lines were significantly more sensitive to carbon ions compared to γ-rays per unit physical dose (Gray/Gy), consistent with previous studies^30,31^. The surviving fraction at 2 Gy (SF2) to γ-rays ranged from 0.51-0.78 and at 3.5 Gy (SF3.5) from 0.20-0.51, indicating a broad range of inherent radioresponses. Alternatively, the SF2 values for carbon ions ranged from 0.09-0.23 and SF3.5 from 0.01-0.05, indicating much less cell specific variability. Relative biological effectiveness (RBE) using 10% SF was estimated to be 1.96-3.04 and 2.38-3.53 when using the Mean Inactivation Dose (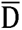). The more radioresistant cell lines to CIRT were PANC.03.27 and PANC-1 and the most radiosensitive cell lines were CAPAN-1 and PANC.04.03 (**Fig. 1A and B**).

**Figure 1.**
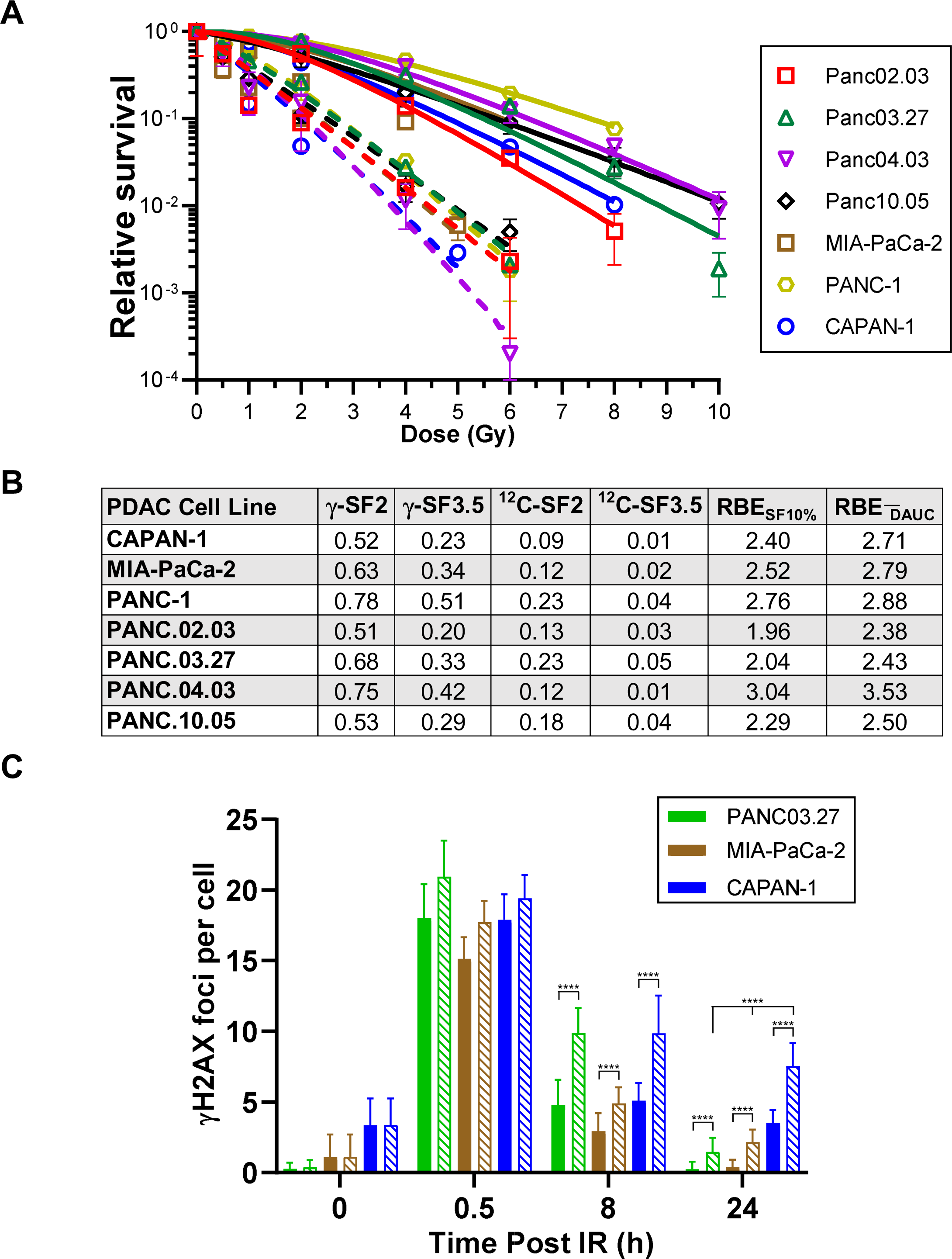
**A**. Clonogenic survival assays were performed to compare the radiation sensitivities of seven PDAC cell lines. Cells were irradiated at the indicated doses of γ-rays (solid) or carbon ions (dashed) and plated for analysis of survival and colony-forming ability. **B.** Survival fraction (SF) at 2 and 3.5 Gy in response to γ-rays and carbon ions. Relative biologic effectiveness (RBE) calculated using multiple methods. RBE_SF10%_, RBE calculated using 10% survival and RBE_DAUC_, RBE calculated using mean inactivation dose derived from Reimann sum. **C.** Immunostaining of γH2AX foci in PANC03.27, MIA PaCA-2, and CAPAN-1 cells after exposure to 1 Gy of γ-rays (solid) or carbon ions (crosshatch). Cells were fixed 0.5, 8, and 24 h after IR and immunostained for γH2AX foci. γH2AX foci were counted for each cell and averaged. Student’s *t*-test (two-sided) was performed to assess statistical significance (\*\*\*\**P* < 0.0001).

The radioresponse to γ-ray and CIRT correlated with unrepaired DNA double strand breaks (DSBs), as γ-H2AX (surrogate marker from DSBs) focus resolution was attenuated in a higher proportion in response to ^12^C ions than γ-rays in the representative cell lines PANC03.27, MIA-PaCa-2, and CAPAN-1 at 8-and 24-hours post-IR (**Fig. 1C**). Moreover, the number of CIRT-induced γ-H2AX foci were significantly higher at 24 hours in the known homologous recombination (HR)-defective cell line, CAPAN-1, compared to PANC.03.27 and MIA-PaCA-2 cells. The PANC.04.03 cell line had the highest RBE among the lines (**Fig. 1A and B**) likely due to its relative radioresistance to γ-rays, suggesting that this radioresistance was diminished when exposed to CIRT. Evaluation of the cBioPortal and COSMIC databases failed to identify a pathogenic mutation or deletion in a DDR gene in the PANC.04.03 cell line, indicating an unknown factor/phenotype is driving the increased sensitivity to CIRT. High γH2AX focus formation in G1 cells without exogenous stress indicates increased intrinsic replication stress^32^; therefore, we assessed if this was increased in PANC.04.03 cells. We found that PANC.04.03 has increased γH2AX foci in G1 cells without exogenous stress, suggesting that this cell line has elevated replication stress (**Fig. 2A**). Moreover, increased DNA damage in the PANC.04.03 cell line in the absence of treatment with an exogenous DNA damaging agent was found via immunoblotting which shows that DNA-PKcs is autophosphorylated at serine 2056 and KAP1 and CHK2 are phosphorylated at serine 824 and threonine 68, respectively, in untreated cells (**Fig. 2B, compare UT lanes**). Finally, cells with increased replication stress are sensitive to ATR inhibition^26^. We found that PANC.04.03 cells are more sensitive to ATRi treatment than PANC.03.27 and MIA-PaCa-2, which further supported the notion that PANC.04.03 have elevated replication stress (**Fig. 2C**). Collectively, our data indicates that HR defects and/or unstable genomes via elevated replication stress in PDAC cells may affect the cellular response to CIRT.

**Figure 2.**
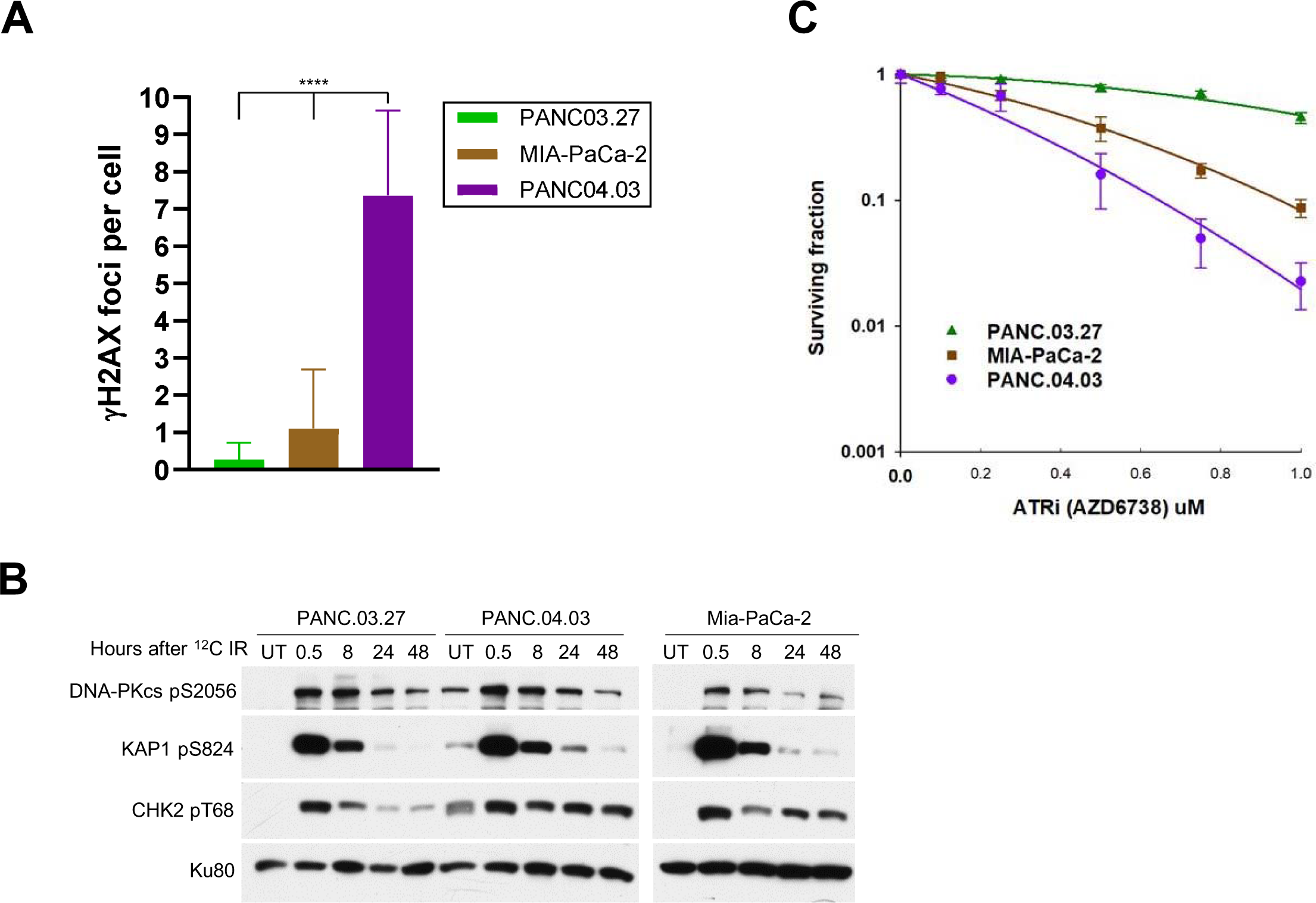
**A**. Immunostaining of γH2AX foci in PANC03.27, MIA-PaCA-2, and PANC.04.03 in G1 cells in the absence of exogenous DNA damage. γH2AX foci were counted for each cell and averaged. Student’s *t*-test (two-sided) was performed to assess statistical significance (\*\*\*\**P* < 0.0001). **B.** PANC03.27, MIA-PaCA-2, and PANC.04.03 were irradiated using a dose of 0 (Untreated, UT) or 1 Gy carbon ions and then allowed to recover at times indicated in the figure. Phosphorylation of DNA-PKcs at S2056, KAP1 at S824, and CHK2 at T68 were assessed via immunoblotting. **C.** Clonogenic survival assays were performed to compare the sensitivities of PANC03.27, MIA-PaCA-2, and PANC.04.03 to increasing doses of the ATR inhibitor AZD6738.

### Non-homologous end joining and homologous recombination mediate the repair of ^12^C Ion induced DSBs

The pathway or pathways integral to the repair of DSBs induced by therapy-relevant carbon ions in PDAC cell lines is still not clear. Thus, we aimed to determine if a specific DSB repair pathway is required for the repair of γ-ray and carbon ion induced damage in PDAC cell lines. Specifically, we pretreated cells with either the DNA-PK_cs_ inhibitor NU7441, RAD51 inhibitor B02, or ATR inhibitor AZD6738 to inhibit the NHEJ, HR, and ATR-CHK1 pathways, respectively. Clonogenic survival assays show that inhibiting the NHEJ pathway resulted in a marked sensitivity to γ-rays in the PANC.03.27 (**Fig. 3A**) and MIA-PaCa2 (**Fig. 3C**) cells. No significant increase in radiosensitivity to γ-rays was observed when cells were pretreated with the RAD51 and ATR inhibitors (**Figs. 3A and C**). This observation correlated with unrepaired DNA double strand breaks (DSBs), as treatment with the DNA-PK_cs_ inhibitor resulted in a significant increase in unrepaired DSBs as monitored by γH2AX foci resolution at 8– and 24-hours post-IR (**Fig. 3B and D**). Next, we assessed clonogenic survival in response to CIRT with the three inhibitors. As shown in **Fig. 3A and C**, treatment with either the DNA-PK_cs_, RAD51, or ATR inhibitor resulted in increased radiosensitivity to ^12^C ions, with significant cell killing in response to pretreatment with the DNA-PK_cs_ and RAD51 inhibitors. Similar to the γ-ray data, the increased radiosensitivity correlated with unrepaired DSBs (**Fig. 3B and D**). Moreover, we assessed if ATM inhibition affected the response to CIRT. We found that pretreatment with the ATM inhibitor KU55933 resulted in a marked radiosensitization in PANC.03.27, MIA-PaCa-2, and PANC.04.03 (**Supplementary** Fig. 3A-C). We also examined if alt-EJ played a role in response to CIRT-induced DNA damage by examining if pretreatment with an inhibitor to PARP (olaparib) affects radiosensitization to CIRT in the PANC.03.27 cell line. We observed that treatment with olaparib did not result in increased radiosensitization to carbon ions (**Supplementary** Fig. 3A). Finally, we found that inhibition of the protein kinases CDK4/6, which are required for the transition the transition to G1 to S phase, did not affect the cellular response to CIRT (**Supplementary** Fig. 3B). Collectively, the data support that the multiple DDR proteins and pathways are required for the repair of carbon ion generated DSBs, including NHEJ, HR, ATR, and ATM.

**Figure 3.**
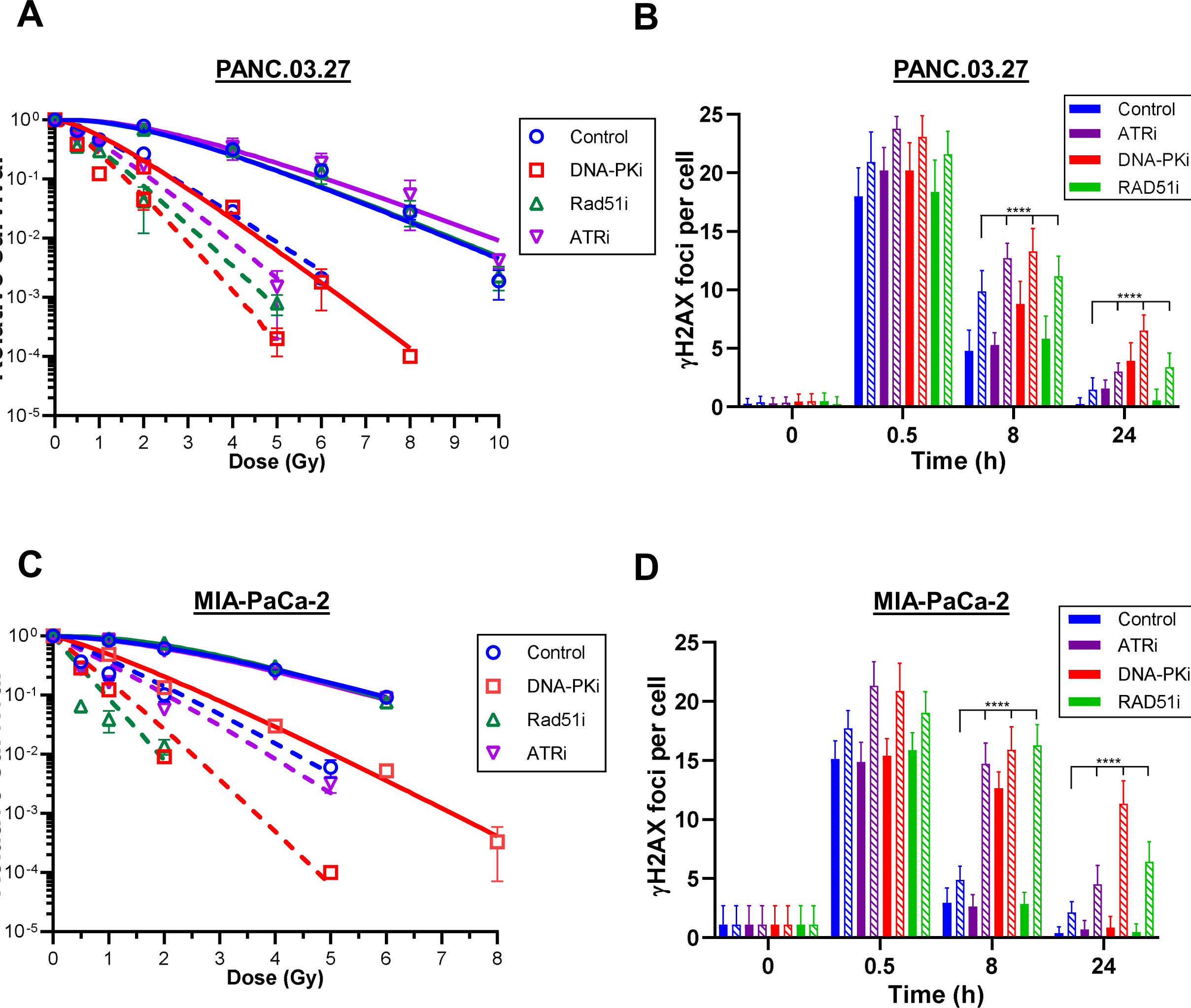
Clonogenic survival assays were performed to compare the radiation sensitivities of PANC03.27 (**A**) and MIA-PaCa-2 (**C**) cells in the presence of DNA damage response inhibitors. Cells were preteated with DMSO (Control), 3 μM NU7441 (DNA-PKi), 20 μM B02 (RAD51i), or 100 nM AZD6738 (ATRi) and then irradiated at the indicated doses of γ-rays (solid) or carbon ions (dashed) and plated for analysis of survival and colony-forming ability. Immunostaining of γH2AX foci in PANC03.27 (**B**) and MIA PaCA-2 (**D**) cells after pretreatment with DMSO (control), 3 μM NU7441 (DNA-PKi), 20 μM B02 (RAD51i), or 100 nM AZD6738 (ATRi), and then exposed to 1 Gy of γ-rays (solid) or carbon ions (crosshatch). Cells were fixed 0.5, 8, and 24 h after IR and immunostained for γH2AX foci. γH2AX foci were counted for each cell and averaged. Student’s *t*-test (two-sided) was performed to assess statistical significance (\*\*\*\**P* < 0.0001).

### Inhibition of HR is a more viable option as a radiosensitizer compared to NHEJ inhibition

Our data indicate that treatment with DNA-PK_cs_, RAD51, and ATR inhibitors mediates increased radiosensitivity to CIRT, suggesting that each of these could potentially be used as radiosensitizers. However, a common issue with radiosensitizers is that they not only sensitize the tumors, but also the normal tissue in the IR field, which can result in adverse tissue events^33^. To assess this, we performed clonogenic survival assays in the entry portion of the depth-dose distribution, which corresponds to the portion of the carbon ion beam expected to predominantly traverse normal tissue. The LET at this entrance position (15 mm) correlates with a LET of 13.0-16.4 keV/µm (**Fig. 4A**). Survival assays found that the radiation response at the entrance position of the carbon ion beam was similar to γ-rays, with the RBE_SF10%_ being 1.08 (**Fig. 4B and C**). Similar to the γ-ray data, pretreatment with either RAD51 or ATR inhibitors did not result in an increase in radiosensitivity at the entrance position of the carbon ion beam, but addition of the DNA-PK_cs_ inhibitor resulted in a significant increase in radiosensitivity (**Fig. 4B**). Taken together, the data suggest that DNA-PK_cs_ inhibitors are not a viable option as a clinical radiosensitizer when combined with carbon ions given the potential for normal tissue toxicity and that inhibitors of HR or ATR are better choices.

**Figure 4.**
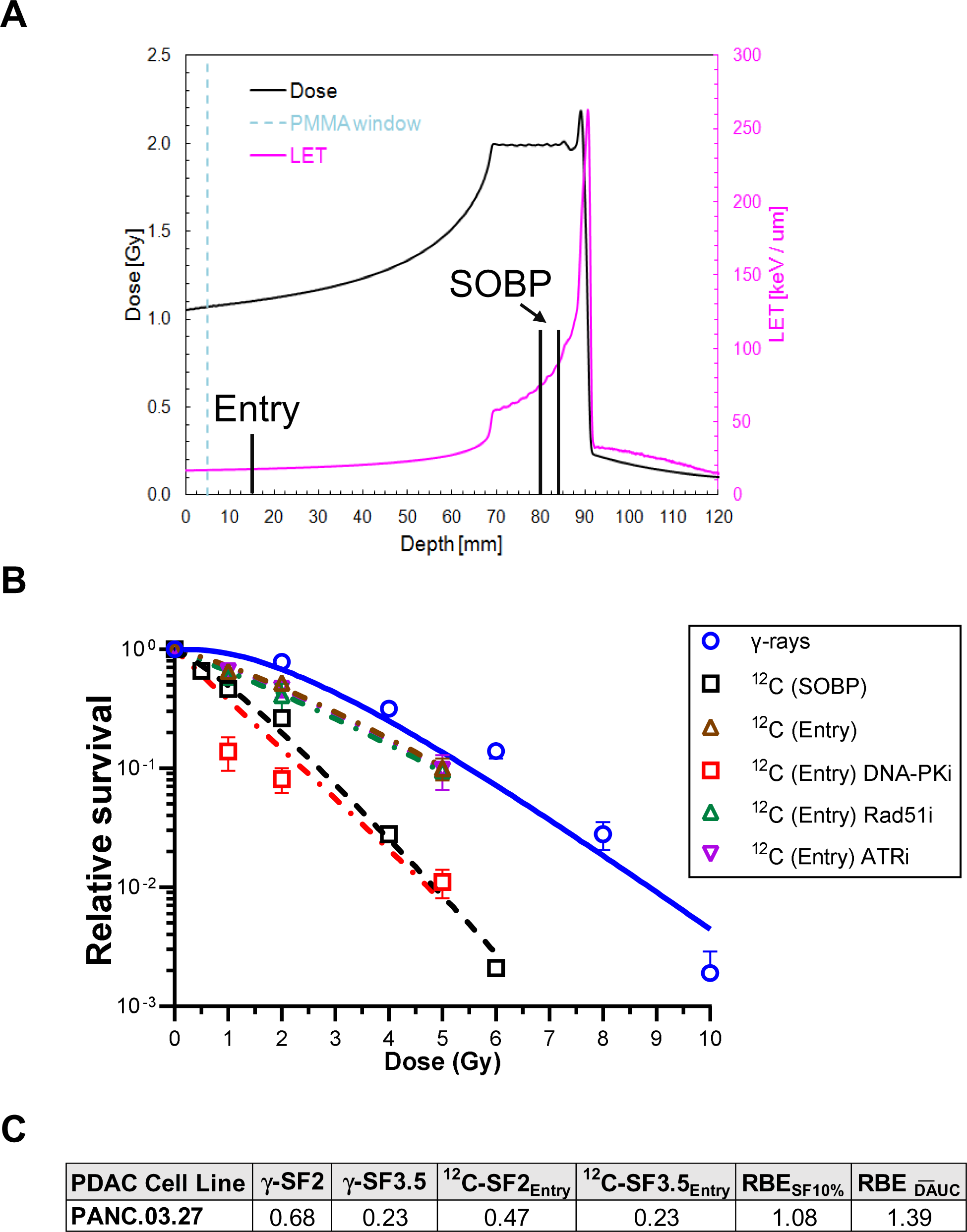
**A**. Schematic showing the placement of the cells at the entry and spread-out Bragg peak (SOBP). **B.** Clonogenic survival assays were performed to compare the radiation sensitivities of PANC03.27 and MIA-PaCa-2 cells in the presence of DNA damage response inhibitors when exposed to carbon ions at the beam entry. Cells were preteated with DMSO (Control), 3 μM NU7441 (DNA-PKi), 20 μM B02 (RAD51i), or 100 nM AZD6738 (ATRi) and then irradiated at the indicated doses of γ-rays (solid) or carbon ions (dashed) and plated for analysis of survival and colony-forming ability. **C.** Survival fraction (SF) at 2 and 3.5 Gy in response to γ-rays and ^12^C ions. Relative biologic effectiveness (RBE) calculated using multiple methods. RBE_SF10%_, RBE calculated using 10% survival and RBE_DAUC_, RBE calculated using mean inactivation dose derived from Reimann sum.

### Treatment with carbon ions results in a differential expression of genes as compared to **γ**-rays

The transcriptomic response of PDAC cells with carbon ions compared to that of γ-rays has not been fully examined. To assess this, we treated five PDAC cell lines (MIA-PaCa-2, PANC.03.27, CAPAN-1, PANC.04.03, and PANC.02.03) with carbon ions or γ-rays and performed expression analysis 2-, 8-, and 24-hours post-treatment against untreated cells at the same times post-irradiation. Per unit dose, transcriptome analysis found that CIRT induces a greater transcriptomic change in pancreatic cancer cell lines than γ-rays. This is demonstrated by the larger number of genes differentially expressed in response to CIRT (557 genes) compared to γ-rays (89 genes) (**Fig. 5A**) and greater variance in principal components for carbon-irradiated cells than the γ-ray-irradiated samples when both are compared against the unirradiated samples using supervised principal component analysis (PCA) (**Fig. 5B**). The signaling pathways most highly enriched with differentially expressed genes in response to radiation were similar between the carbon and γ-ray groups, as was their activation/suppression status. However, carbon particle irradiation induced the expression of several signaling pathways that were not identified in the γ-ray response based upon z score determination. These include the upregulated pathways, “Role of CHK Proteins in Cell Cycle Checkpoint Control”, “p53 Signaling”, “Role of BRCA1 in DNA Damage Response”, and “Ferroptosis Signaling Pathway”, and the downregulated pathway “SPINK1 General Cancer Pathway”. These pathways illustrate an increased DDR, with the activation of pathways that promote HR more prominent in the carbon ion irradiated cell lines per unit dose. Collectively, our data indicates PDAC tumors with defects in HR and/or those with high inherent replication stress respond to CIRT without the concern for excessive normal tissue toxicity.

**Figure. 5.**
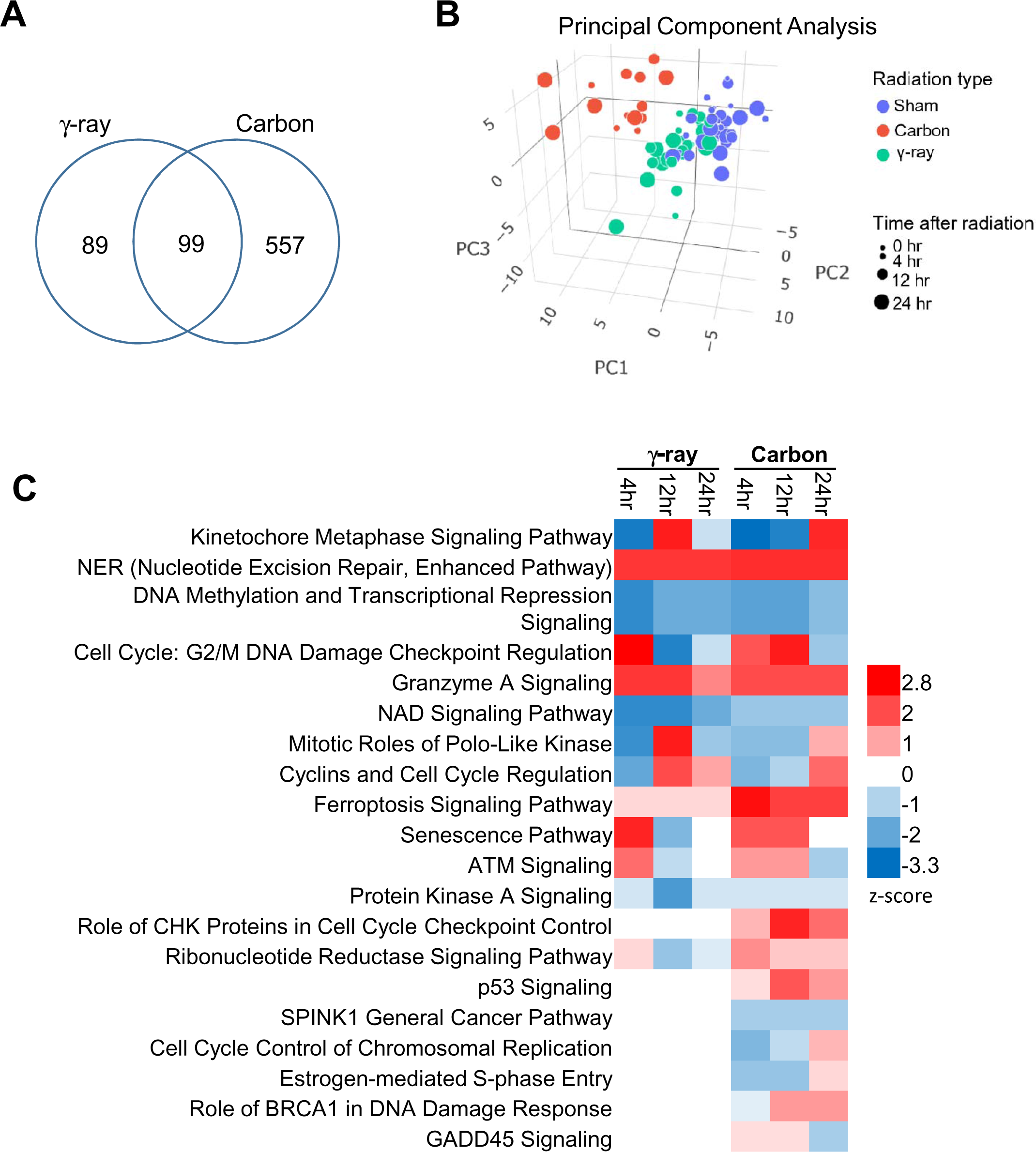
Gene expression analysis of pancreatic cancer cell lines irradiated with γ-rays or carbon particles. **A.** Venn-diagram showing numbers of differentially expressed genes after γ-ray or carbon ion irradiations in comparison to sham-irradiated cell lines (FDR < 0.1). **B.** Plot of cell line samples with the top 3 principal components after supervised PCA analysis using 745 differentially expressed genes. **C.** Pathway analysis of differentially expressed genes in response to radiations showed significantly changed signaling pathways (FDR < 0.1 in at least one condition) with activation z-scores. A positive z-score indicated up-regulation and a negative z-score indicated down-regulation of the pathway.

## DISCUSSION

The proposed benefits from carbon ions in the treatment of malignant tumors in general, and PDAC in particular, is mostly motivated by the physical properties of high-LET radiation beams allowing for the delivery of a more conformal dose distribution to the target as well as a greater biological effect per unit dose, which is described by the RBE. The RBE is strongly influenced by both physical and biological factors and the endpoint chosen. In this study, the RBE of carbon ions was primarily determined by *in vitro* clonogenic cell survival experiments where established cell lines were exposed to various doses of high LET carbon ions as expected in a typical Bragg peak. Here, we observe that the RBE range is 1.96-3.04 (SF10%) and 2.38-3.53 (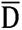) for PDAC cell lines with the clinical beam at CNAO.

Currently, there is little opportunity to optimize CIRT or CIRT plus targeted agents for a given individual based upon their known or determined radiosensitivity and/or targeted vulnerabilities as identified by genetics/genomics. We believe that the genetics of the tumor play a significant role, as defects in the DNA damage response impact the RBE and the tumor response to CIRT. Our data shows that PDAC cell lines with defects in HR and increased replication stress are more sensitive to CIRT than those without. If possible, the genetics/genomics of the patient’s tumor should be taken into account when choosing between therapeutic options. We hypothesized that there would be differential DNA repair pathway(s) required for the response to clinical ion beams in human PDAC cell lines. However, our findings suggest this differential was most profound for γ-rays that are dependent upon the DNA-PK_cs_-dependent NHEJ pathway. In CIRT, we observed the greatest sensitivity for DSBs was with the combination therapy of DNA-PK_cs_-dependent NHEJ pathway and RAD51 HR pathway inhibitors. Furthermore, our results suggest that the ATR pathway is involved in the repair of DSBs induced by clinical carbon ion beams not observed with γ-rays. These results are similar to those obtained with NHEJ and HR-defective Chinese hamster ovary (CHO) cell lines ^20^ and human skin fibroblasts treated with a DNA-PKi ^34^. In addition, we found that PARP inhibition does not affect cell killing by CIRT. Previous work showed that combining CIRT with the PARPi olaparib enhanced radiosensivity in the BRCA1-mutated triple-negative breast cancer cell line HCC1937, but had no effect on the BRCA wild-type cell line MDA-MB-231^35^. We postulate that PARP activity is not required for the response to CIRT-induced DNA damage unless the cell line is HR-defective. Finally, our data indicates that HR-defective cancer cells can be effectively targeted by CIRT. This is of importance as approximately 10% of PDAC tumors are HR-defective and approximately 20% have a defect in the DNA damage response ^23,26,36,37^. Our preclinical data extends this to the use of inhibitors of the HR pathway to potentiate CIRT, where pretreatment with RAD51i, B02, results in increased CIRT-induced cell killing *in vitro* and *in vivo*. A similar effect was seen for proton exposures of lung cancer cells defective in HR ^38^, although the RBE difference was not as great as that seen with CIRT. Additionally, B02 treatment did not increase cell killing at the entry of the depth-dose distribution, representative of normal tissue, suggesting that normal tissue will not be greatly affected by inhibiting HR. However, it should be noted that no HR inhibitor has been approved by the FDA yet and thus there is a need for effective clinical application of an HR inhibitor with favorable properties.

In conclusion, we evaluated CIRT on a cohort of PDAC cells lines and found that the cell lines with defects in HR and those that have increased replication stress are exquisitely more sensitive to CIRT than γ-rays. Further, we elucidated the molecular mechanisms underlying the effects of CIRT on PDAC cell lines and found that NHEJ and HR, but not alt-EJ, are required for the repair of carbon ion induced DSBs. Moreover, we found that targeting a radioresistant PDAC cell line with the HR inhibitor B02 results in markedly increased sensitivity to CIRT at the SOBP but not as effectively at the entry of the depth-dose distribution in *vitro*. Lastly, while agents that target the HR pathway demonstrated no enhanced TGD of PDAC tumors *in vivo*, further experiments would be appropriate to combine HR inhibitors with CIRT *in vivo* to further test this hypothesis, especially where no DDR defect has been identified.

## MATERIALS AND METHODS

### Cell Culture

The cell lines PANC02.03, PANC03.27, PANC04.03, PANC10.05, MIA-PaCa-2, PANC-1, and CAPAN-1 were purchased from American Type Culture Collection (ATCC). PANC02.03, PANC03.27, PANC04.03, PANC10.05 were cultured in RPMI-1640 Medium supplemented with 10 Units/mL human recombinant insulin and 15% fetal bovine serum (FBS). MIA-PaCa-2 and PANC-1 were cultured in Dulbecco’s modified Eagle’s medium supplemented with 10% FBS. CAPAN-1 were cultured in Iscove’s Modified Dulbecco’s Medium supplemented with 20% FBS. The cells were grown in an atmosphere of 5% CO_2_ at 37°C. To inhibit the activity of DNA-PK_cs_, ATR, Rad51 nucleofilament formation, ATM, PARP1, or CDK4/6, cells were incubated for 2 h before the experimental with 3 μM NU7441 (SelleckChem), 100 nM AZD6738 (SelleckChem), 20 μM B02, (SelleckChem), 3 μM KU55933 (SelleckChem), 1 μM olaparib (SelleckChem), or 1 μM palbociclib, respectively.

### Photon Irradiations

Photon irradiation was conducted at the University of Texas Southwestern Medical Center (UTSW) using a J. L. Shepherd sealed horizontal ^137^Cs-sourced irradiator. Dosimetry for this sealed source irradiators is validated on an annual basis. Briefly, cells in culture were placed on a 360° platform revolving at 13 RPM, irradiated, removed from the irradiator, and immediately returned to the incubator. At the time of treatment, the irradiator had a dose rate of ∼3.25 Gy/min.

### ^12^C Ion Irradiations

All ^12^C ion irradiations were performed at the Centro Nazionale di Adroterapia Oncologica (CNAO) facility in Pavia, Italy utilizing established methods ^39^. Specifically, cells were irradiated in T12.5 cm flasks while immersed in a water bath at 37°C using CNAO’s clinical pencil beam scanning ^12^C-ion beam. A spread-out Bragg peak (SOBP) was created to deliver a homogenous (± 2.5%) physical dose across the target volume. The beam quality has been previously characterized ^40^ and adheres to the recommendations of a NCI special panel on particle beam characterization ^41^. The dimensions of the SOBP were 17 cm in width, 7 cm in height, and 2 cm in depth (**Sup.** Fig. 2B). Cells were centered in the SOBP using a leucite holder with the adherent cells aligned in their flasks back-to-back such that the depth of the cells in the upstream flask was 80.0 mm water equivalent depth (WED) while the position of cells in the downstream flask was 84.0 mm of WED (**Sup.** Fig. 2A). LETs at the aligned positions were 74.1 and 89.3 keV/µm, respectively at a typical dose rate of 0.60 Gy/min. (No difference in biological response was observed based upon position.) The entrance LET was 13.0-16.4 keV/µm at a depth of 15 mm.

### Clonogenic Cell Survival Assays

Cells undergoing log phase growth at roughly 70%–80% maximum cell culture density were trypsinized and then seeded into T-12.5 flasks at low density in complete growth medium 8 h prior to irradiation. If using an inhibitor, the regular media was changed and supplemented with the specific inhibitor and allowed to incubate for 2 hr. Five minutes prior to irradiation with either γ-rays or carbon ions, cell culture flasks were filled to the neck with complete growth media containing 2% FBS with or without inhibitors. Cells were irradiated with doses of 1, 2, 4, 6, and 8 Gy of γ-rays, or 0.5, 1, 2, and 4, 5, and/or 6 Gy carbon ions. Following irradiation, growth medium containing 2% FBS was immediately aspirated and replaced with growth medium containing 10% FBS and dishes were allowed to incubate for ∼10 population doublings based on cell-specific doubling times. Media with 2% FBS was used during the irradiations to limit the overall volume of FBS needed to completely fill the T12.5 flasks for only a few minutes as was necessary at CNAO. Using 2% FBs for such a limited time had no effect on cell growth nor radioresponse. Following incubation, the media was removed and the cultures were incubated in 100% ethanol solution containing 0.1% crystal violet in order to fix and stain the colonies. After drying, colonies were counted to determine the number of surviving cells following irradiation. Only colonies identified as having more than 50 cells per colony were scored as surviving, and the surviving fraction was determined by dividing the number of colonies by the product of the plating efficiency of the cell line multiplied by the number of cells seeded.

### Survival Curve Fits

Survival curves were fitted based upon the Repairable Conditionally Repairable (RCR) Model as described in Equation 1 where d is the dose per fraction and a, b, and c are parameters determined using a curve fitting algorithm ^42^.

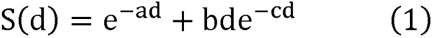

The γ-ray survival assays were performed at least twice for each cell line. If the coefficient of variation at 2 Gy was greater than 25%, they were repeated.

### RBE Calculations

Because all experiments were performed as single exposure irradiations the RBE values were calculated by using the simplest determination of RBE, i.e., by comparing a radiosensitivity value from ^137^Cs exposures (reference) to that same radiosensitivity value determined from ^12^C exposures (test) as in Equation 2.

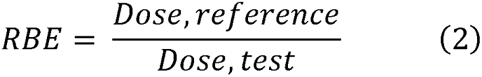

Radiosensitivity parameters included:

*Dose at 10% survival:* The dose at SF_10%_ was calculated using values generated with the RCR model as described in Equation 1.

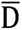: 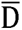 was defined by Kelleher and Hug to represent cell survival across the entire dose response ^43,44^. 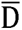 is calculated by 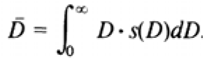 where s(D) = –dS(D)/dD and S(D) is the survival function representing the fraction of cells surviving a radiation dose D. S(D) is the probability that a dose greater than D is required to kill a randomly selected cell. s(D) = –dS(D)/dD defines the corresponding probability density. 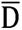 can be represented by the area under the survival curve which in this case was calculated using a trapezoidal method of the area under the curve for each survival assay. 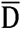 is equal to D_0_ for exponential survival curves.

### **γ**H2AX Foci Kinetics

IR-induced γH2AX kinetics were determined as previously outlined with modifications^45,46^. Cells were grown on coverslips one day before the experiment and on the day of the experiment, cells were exposed to 1 Gy of γ-rays or ^12^C ion using the protocols outlined above. At 0, 0.5, 8, or 24 hr time points after IR, cells were washed twice with ice cold 1× phosphate buffered saline (PBS) and fixed with 4% paraformaldehyde (in 1X PBS) for 20 min at RT, washed 5 times with 1X PBS, and incubated in 0.5% Triton X-100 on ice for 10 min. Cells were washed 5 times with 1X PBS and incubated in blocking solution (5% goat serum (Jackson Immuno Research) in 1X PBS) overnight. The blocking solution was then replaced with the γH2AX (Cell Signaling Technology, 7631) and/or 53BP1 (ab175933, Abcam) primary antibody diluted in 5% goat serum in 1X PBS and the cells were incubated for 2 h. Cells were then washed 5 times with wash buffer (1% BSA in 1X PBS). Cells were incubated with the Alexa Fluor 488 (Molecular Probes) and/or anti-mouse IgG conjugated with Texas Red (Molecular Probes) (1:1000 dilution for both antibodies) secondary antibodies in 1% BSA, 2.5% goat serum in 1X PBS for 1 h in the dark, followed by five washes. After the last wash, cells were mounted in VectaShield mounting medium containing 4’6-diamidino-2-phenylindole (DAPI). The images were acquired using a Zeiss Axio Imager fluorescence microscope utilizing a 63X oil objective. ≥50 cells were analyzed for each time point.

### Isolation of RNA from Pancreatic Cancer Cell Lines

Pancreatic cancer cell lines were irradiated with 1 Gy of γ-rays (UTSW) or carbon particles (CNAO). Total RNA was isolated at 4-, 12-, and 24-hours post-irradiation. Total RNA from sham-irradiated cell lines were isolated at the same time as the irradiated samples. The sham-irradiated cells were also collected right before irradiation and used as baseline (0 hour) references. RNA extractions were performed using miRNeasy Mini Kit (Qiagen) according to the manufacturer’s protocol. Once isolated, RNA concentration was determined using a Nanodrop 2000 Spectrophotometer (Thermo Scientific) and quality was assessed using an Experion Electrophoresis system (Bio-Rad).

### Transcriptomic Analysis of PDAC Cell Lines Following **γ**-ray and ^12^C Ion Irradiations

Sequencing library was prepared using the Illumina (San Diego, CA) Truseq Stranded Total RNA Prep Kit. Paired end next generation sequencing was performed by DNAlink (San Diego, CA) using an Illumina Novaseq 6000 sequencer. Sequencing reads were trimmed to remove adaptor sequences and were aligned with human genome GRCh38 using STAR with the 2-pass option. The aligned reads were quantified using the RSEM program. Gene counts were summarized using tximport ^47^ R library and subsequently normalized with TMM algorithm using edgeR. Differential expression analysis was performed using log2 counts per million (cpm) values and the limma package ^48^. Genes with low read counts were removed by filterByExpr function within the edgeR package using default parameters ^49^. Differential expression analysis was performed by fitting the log2 cpm values using the limma-trend approach described in the R limma package. Differentially expressed genes in irradiated cells at any of the 3 time-points were identified by performing a moderated F-test using FDR < 0.1 as a statistical cutoff. Prior to principal component analysis (PCA), the cell line-specific variances were removed using the limma function and PCA was performed using the R prcomp function. Pathway analysis of differentially expressed genes was performed using the Ingenuity Pathway Analysis software (Qiagen).

## FIGURE LEGENDS

**Supplementary Fig. 1.**
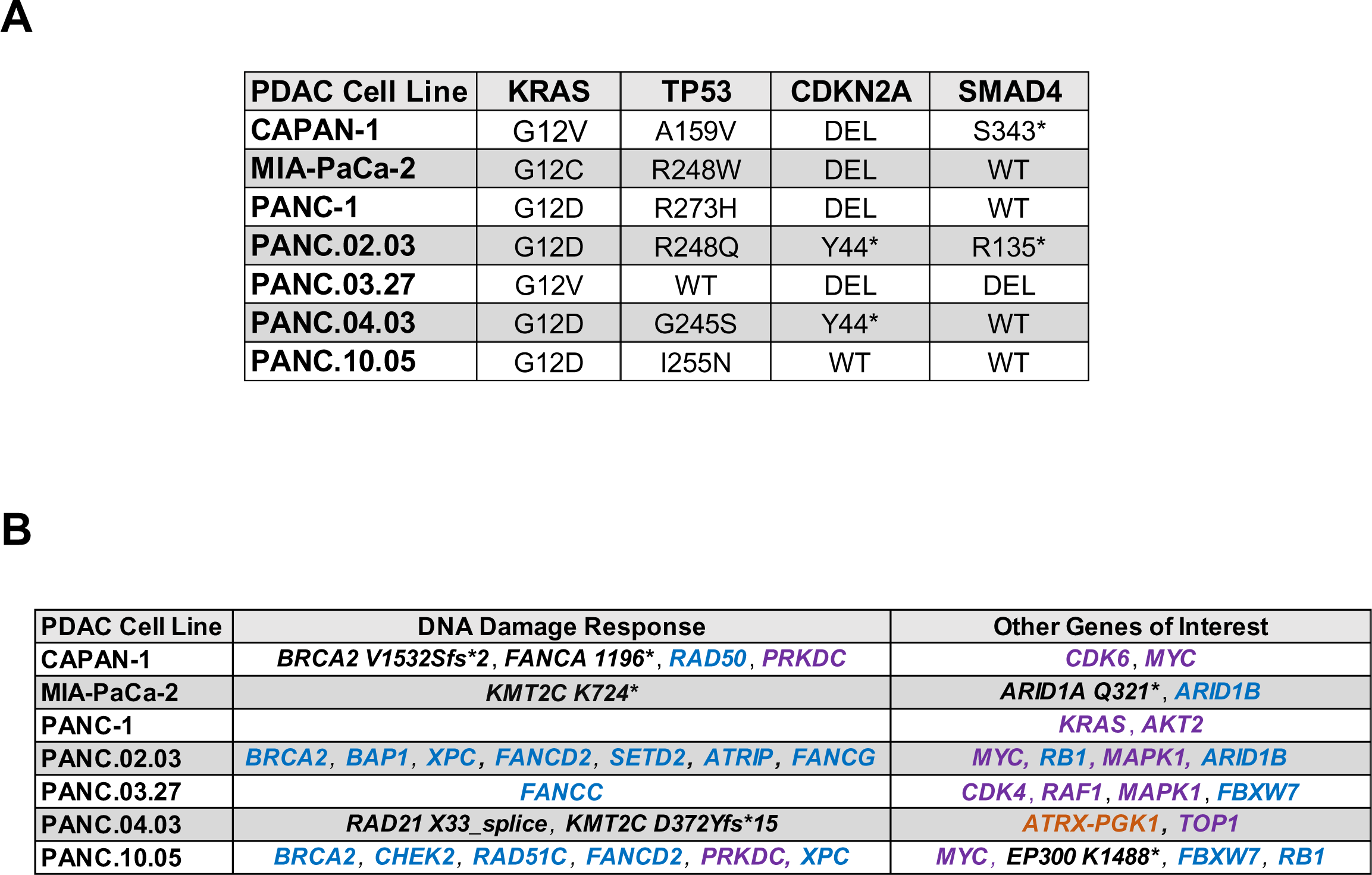
**A**. List of PDAC cell lines used in the study and mutations of the common driver genes associated with PDAC etiology. **B.** List of DNA damage response gene and other genes of interests with mutations, copy number alteration, or structural variants. Point mutations are in **black**, deletions are in **blue**, amplifications are in **purple**, and structural variants in **dark orange**.

**Supplementary Fig. 2.**
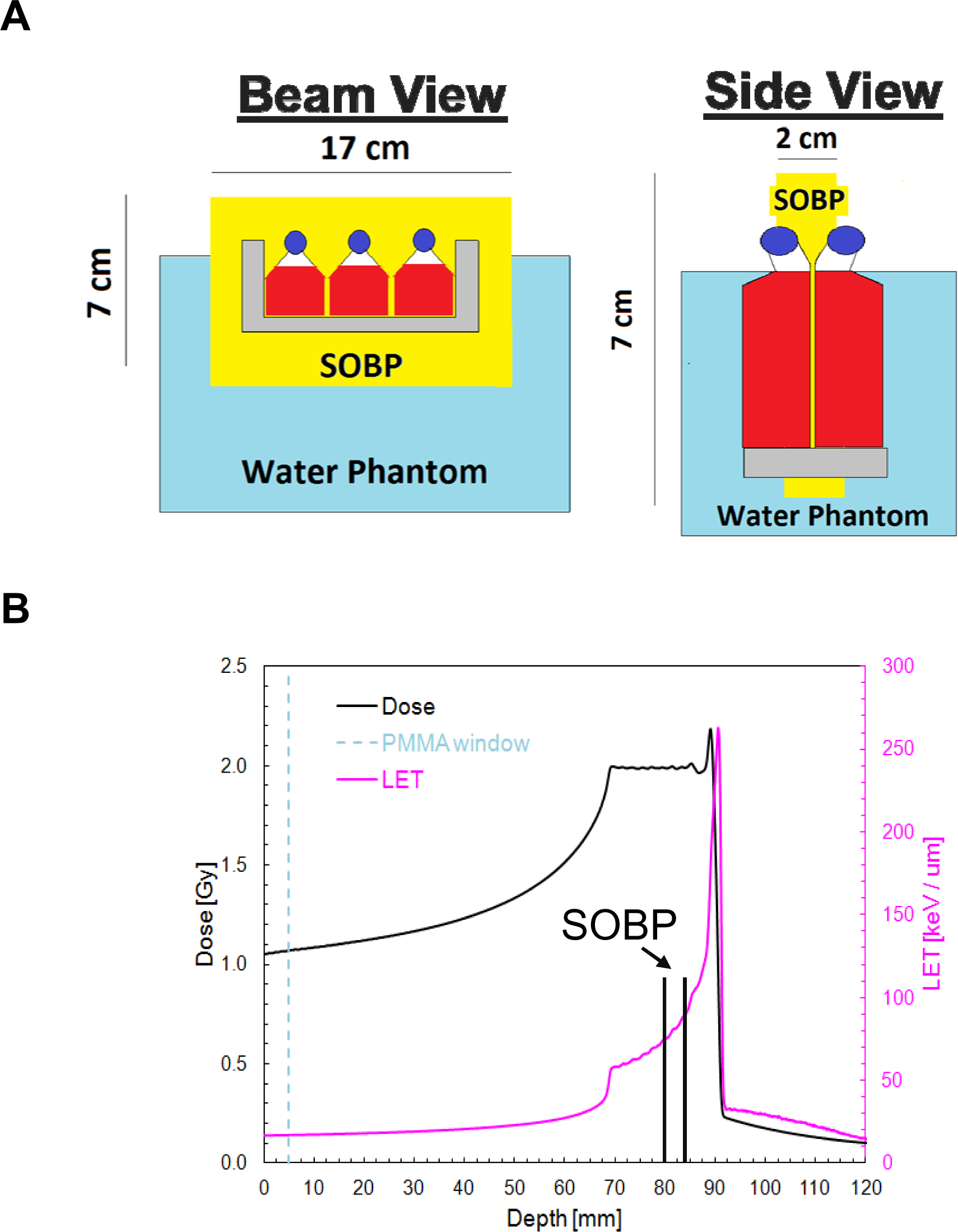
**A**. Schematic representing the SOBP of the carbon ion beam utilized for *in vitro* experiments performed at CNAO. Irradiation occurred in a 7 cm tall, 17 cm wide, and 2 cm deep SOBP with monolayer cell cultures irradiated in t-12.5 culture flasks oriented back to back at the center of the 2 cm SOBP depth. **B.** Relative depth dose curve demonstrating the location of the cell irradiation in the SOBP along the total Bragg curve for carbon ions demonstrating a dose average of LET of ∼ 75 keV/μm.

**Supplementary Fig. 3.**
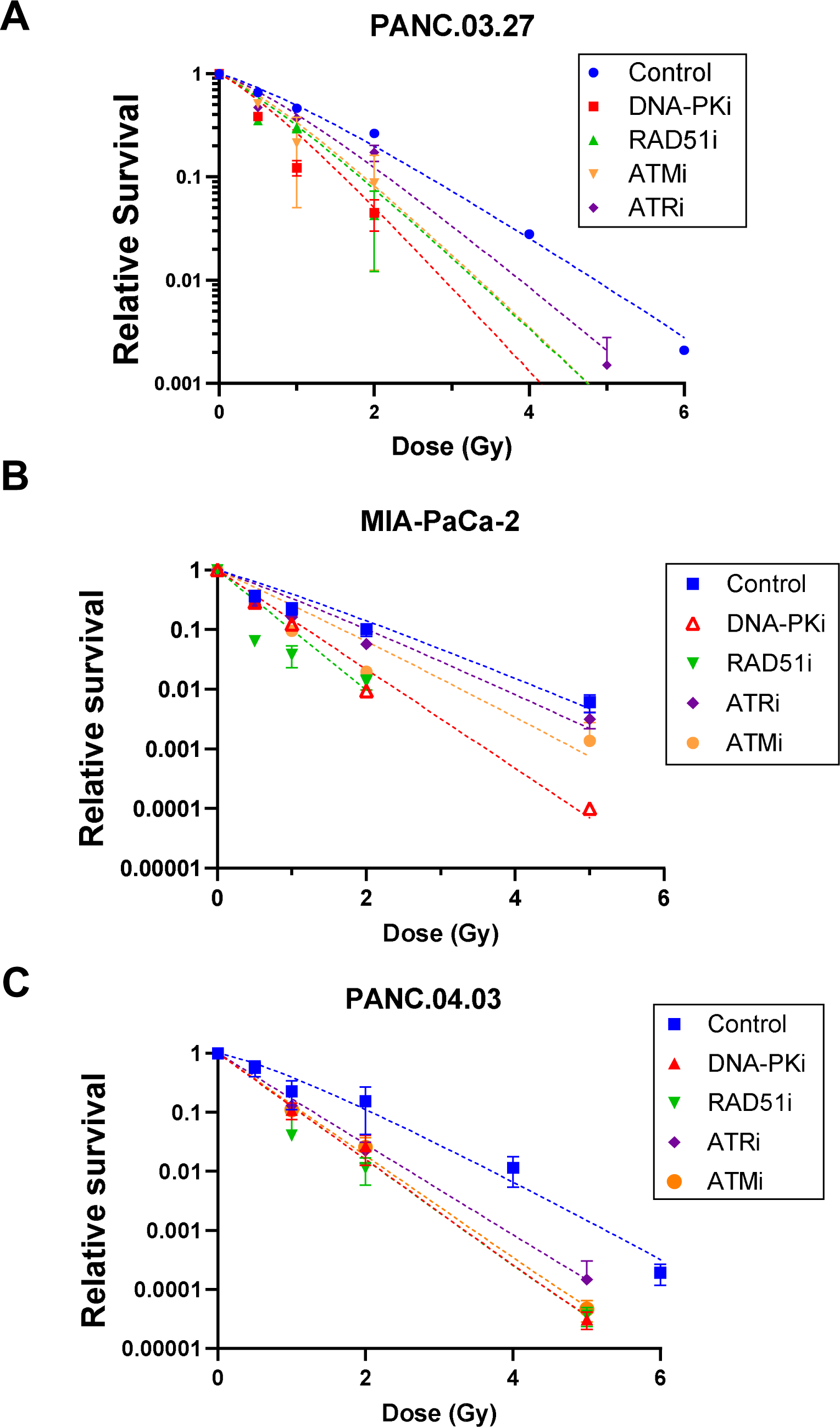
Clonogenic survival curves for the PDAC cell lines **(A)** PANC.03.27, **(B)** MIA-PaCa-2, and **(C)** PANC.04.03 to carbon ions and with the addition of DMSO (Control), NU7441 (DNA-PKi), B02 (RAD51i), AZD6738 (ATRi), or KU55933 (ATMi). Pretreatment with each inhibitor resulted in increased radiosensitization in response to carbon ions in all three PDAC cell lines.

**Supplementary Fig. 4.**
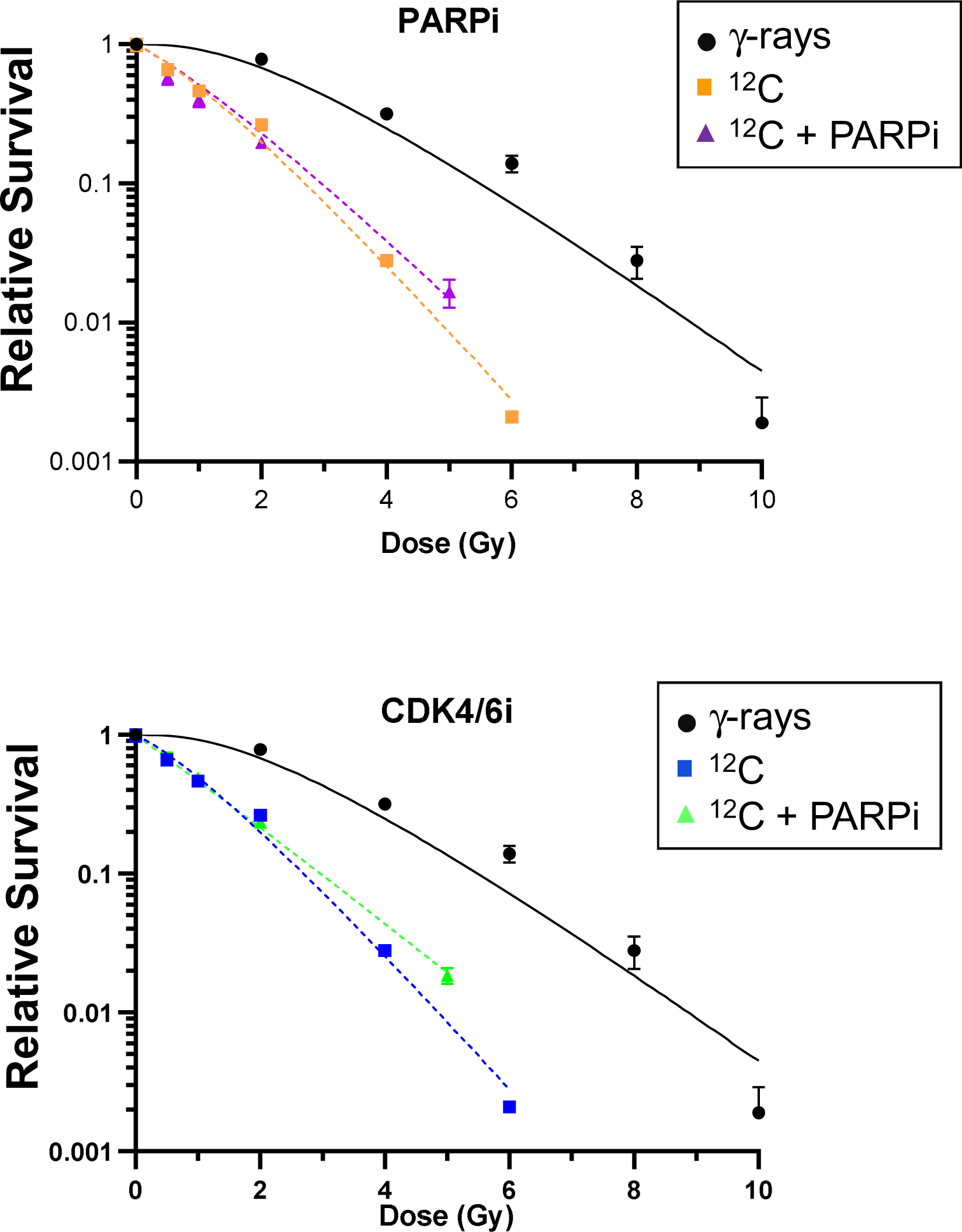
Clonogenic survival curves for the PDAC cell line PANC.03.27 to γ-rays and carbon ions and carbons ions with the addition of the **(A)** PARP inhibitor olaparib or the **(B)** CDK4/6 inhibitor palbociclib. Neither PARPi nor CDK4/6i treatment affected the response to carbon ions.

